# Epigenetic gene silencing by heterochromatin primes fungal resistance

**DOI:** 10.1101/808055

**Authors:** Sito Torres-Garcia, Pauline N. C. B. Audergon, Manu Shukla, Sharon A. White, Alison L. Pidoux, Robin C. Allshire

## Abstract

Genes embedded in H3 lysine 9 methylation (H3K9me)–dependent heterochromatin are transcriptionally silenced^1-3^. In fission yeast, *Schizosaccharomyces pombe*, H3K9me heterochromatin silencing can be transmitted through cell division provided the counteracting demethylase Epe1 is absent^4,5^. It is possible that under certain conditions wild-type cells might utilize heterochromatin heritability to form epimutations, phenotypes mediated by unstable silencing rather than changes in DNA^6,7^. Here we show that resistant heterochromatin-mediated epimutants are formed in response to threshold levels of the external insult caffeine. ChIP-seq analyses of unstable resistant isolates revealed new distinct heterochromatin domains, which in some cases reduce the expression of underlying genes that are known to confer resistance when deleted. Targeting synthetic heterochromatin at implicated loci confirmed that resistance results from heterochromatin-mediated silencing. Our analyses reveal that epigenetic processes allow wild-type fission yeast to adapt to non-favorable environments without altering their genotype. In some isolates, subsequent or co-occurring gene amplification events enhance resistance. Thus, heterochromatin-dependent epimutant formation provides a bet-hedging strategy that allows cells to remain genetically wild-type but transiently adapt to external insults. As unstable caffeine-resistant isolates show cross-resistance to the fungicide clotrimazole it is likely that related heterochromatin-dependent processes contribute to anti-fungal resistance in both plant and human pathogenic fungi.

## Main Text

H3K9me heterochromatin can be copied during replication by a read-write mechanism^4,5^ and has been observed to arise stochastically at various loci, albeit only in the absence of key anti-silencing factors^8-11^. We reasoned that if heterochromatin can redistribute in wild-type *S. pombe* cells epimutations could be generated that allow cells to adapt to external insults. Unlike genetic mutants we predicted that such epimutants would be unstable, resulting in gradual loss of the resistance phenotype following growth in the absence of the external insult. To explore this possibility, we chose to test caffeine resistance because deletion of genes with a wide variety of cellular roles is known to confer resistance^12^, thereby increasing the chance of obtaining epimutations. We also reasoned that such unstable epimutants would occur more frequently at moderate caffeine concentrations that prevent most cells from growing (16 mM) rather than at high stringency selection (20 mM) used in screens for genetic caffeine-resistant mutants^12^.

As other secondary events might also occur upon prolonged growth on caffeine, we froze one aliquot of each isolate as soon as possible after resistant colony formation and then froze consecutive aliquots of each isolate after continued growth on caffeine (Fig. 1a). This provided a time series, permitting detection and separation of potential initiating and subsequent secondary events.

**Figure 1.**
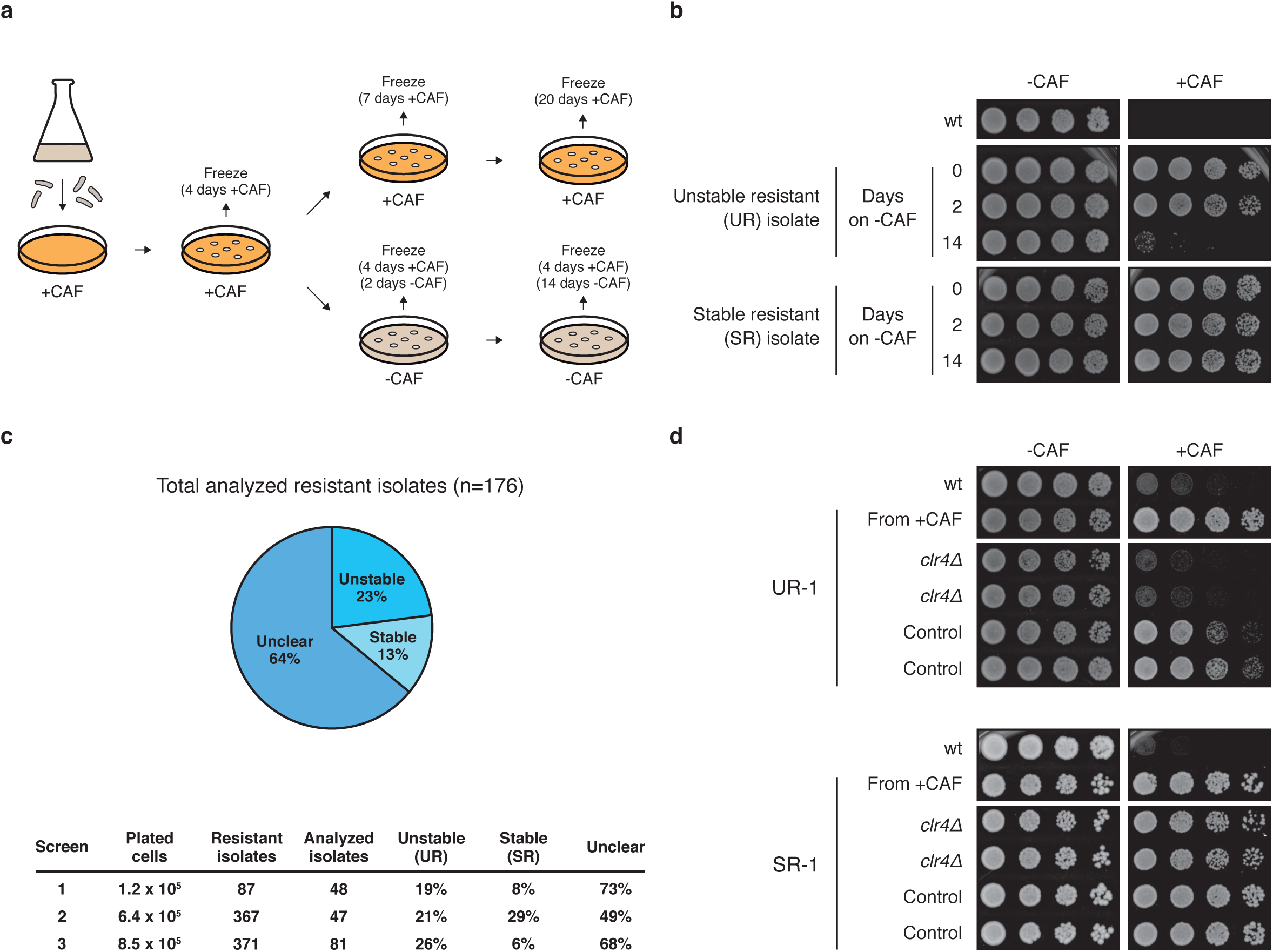
Identification of heterochromatin-dependent epimutants resistant to caffeine. **a**, Schematic of the screening strategy. *S. pombe* wild-type (wt) cells were plated on caffeine-containing (+CAF) plates. Caffeine-resistant isolates were then grown on +CAF plates for 4, 7 or 20 days or on non-selective (-CAF) medium plates for 2 and 14 days. Cells were then serially diluted and spotted on -CAF and +CAF media to assess resistance to caffeine. **b**, Unstable (UR) and stable (SR) caffeine-resistant isolates were identified. After growth on non-selective media for 14 days caffeine resistance is lost in UR isolates but not in SR isolates. **c**, Frequency of unstable (UR) / stable (SR) caffeine-resistant isolates obtained from 3 independent screens. 64% of isolates did not display a clear phenotype (unclear). **d**, Caffeine resistance in UR isolates depends on the Clr4 H3K9 methyltransferase. *clr4*+ (*clr4δ*) or an unlinked intergenic region (Control) were deleted in unstable (UR-1) and stable (SR-1) caffeine-resistant isolates.

Colonies that grew after plating wild-type fission yeast (972 *h*^*-*^) cells in the presence of caffeine (16 mM caffeine, +CAF) were picked. Following freezing, isolates were then successively propagated in the absence of caffeine (-CAF). Re-challenging isolates with caffeine revealed that 23% lost their caffeine resistance after 14 days of non-selective growth (denoted ‘unstable isolates’, UR) whereas 13% remained caffeine resistant (denoted ‘stable isolates’, SR). 64% of isolates did not display a clear phenotype (denoted ‘unclear’) (Fig. 1b, c and Extended Data Fig. 1a, b).

Deletion of *clr4*^+^ encoding the sole H3K9 methyltransferase in *S. pombe*^13,14^ from resistant isolates resulted in immediate loss of caffeine resistance in unstable, but not in stable isolates (Fig. 1d and Extended Data Fig. 1c), indicating that caffeine resistance in unstable isolates requires heterochromatin.

Whole genome sequencing (WGS) of the stable isolate SR-1 uncovered a mutation in *pap1*^+^ responsible for the caffeine-resistant phenotype (Extended Data Fig. 2 and ^15^). ChIP-seq for H3K9me2 on SR-1 revealed no changes in heterochromatin distribution.

WGS of unstable isolates revealed no genetic changes in coding sequences involved in either caffeine resistance or H3K9me2-mediated silencing, and 8 of 30 analyzed unstable isolates had no detectable genetic change compared to wild-type (Supplementary Information Table 1). ChIP-seq for H3K9me2 on unstable isolates revealed an altered heterochromatin distribution (Fig. 2a, b). Unstable resistant isolate UR-1 exhibited a new H3K9me2 domain over the *hba1* locus, whereas UR-2 – UR-6 exhibited H3K9me2 domains over *ncRNA.394, ppr4, grt1, fio1* and *mbx2* loci, respectively (Fig. 2a, b and Supplementary Information Table 1). Deletion of *hba1*^+^ is known to confer caffeine resistance^16^, suggesting that these novel heterochromatin domains may drive caffeine resistance by silencing underlying genes. Accordingly, RT-qPCR analysis revealed reduced expression of genes underlying the observed novel heterochromatin domain at the *hba1* locus (Fig. 2c).

**Figure 2.**
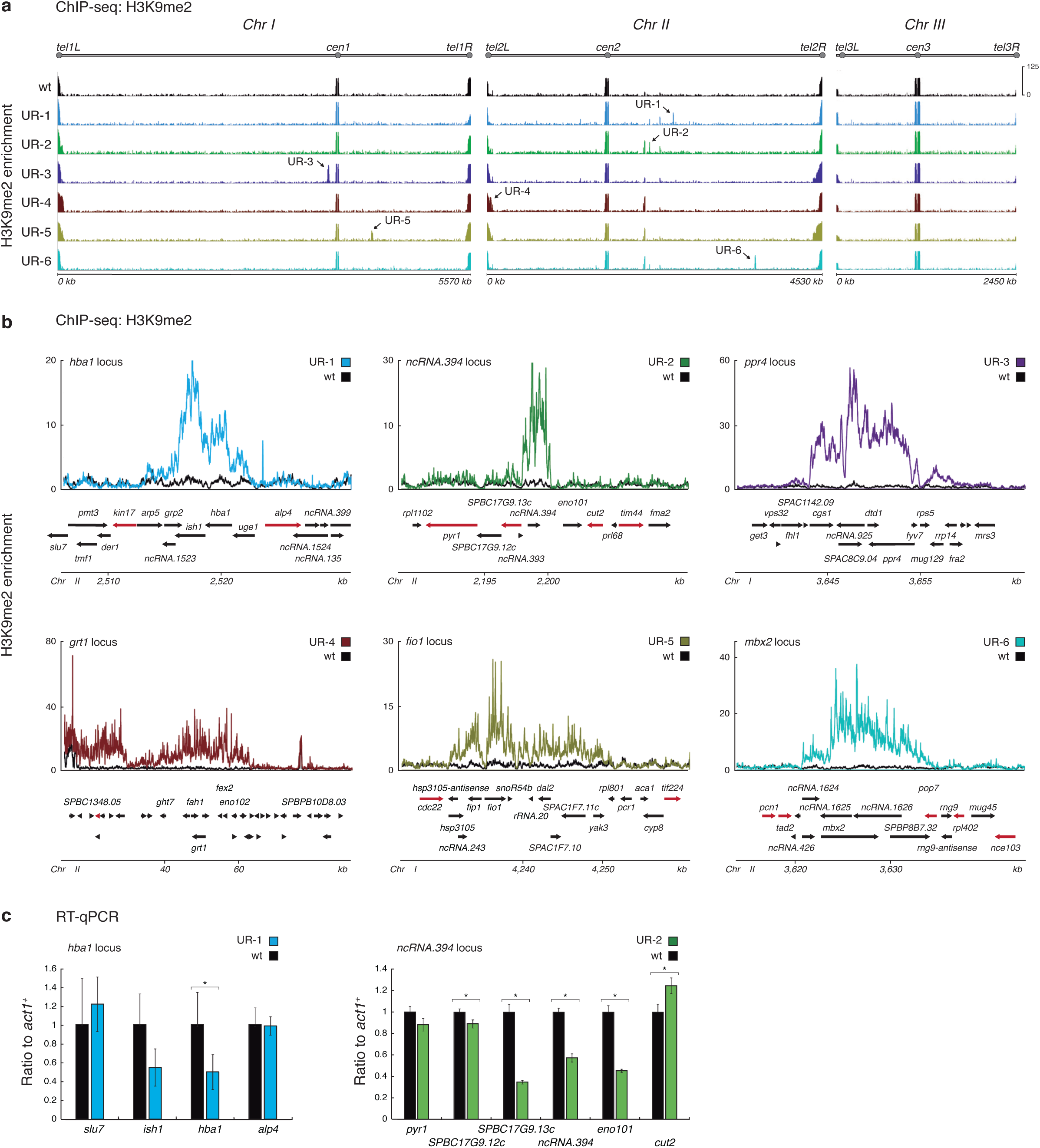
Ectopic domains of heterochromatin are detected in unstable (UR) caffeine-resistant isolatesTorres-Garcia *et al.* **a**, Genome-wide H3K9me2 ChIP-seq enrichment in UR isolates and wt. Data are represented as relative fold enrichment over input. **b**, H3K9me2 ChIP-seq enrichment at ectopic heterochromatin domains in individual isolates. Data are represented as relative fold enrichment over input and compared to levels in wt cells. Relevant genes within and flanking ectopic heterochromatin domains are indicated. Red arrows indicate essential genes. **c**, Gene transcript levels within and flanking ectopic heterochromatin domains in isolates UR-1 and UR-2. Data are mean ± SD (error bars) (n = 3 experimental replicates). * *P* < 0.05 (*t* test).

The *ncRNA.394, ppr4, grt1, fio1* and *mbx2* loci have not previously been implicated in caffeine resistance. Interestingly, 24 of 30 unstable isolates showed an ectopic heterochromatin domain over the *ncRNA.394* locus (Extended Data Fig. 3a and Supplementary Information Table 1), and reduced levels of transcripts were present (Fig. 2c), suggesting that transcriptional silencing within this region might mediate caffeine resistance. *ncRNA.394* was previously described as a heterochromatin ‘island’^8^, yet H3K9me2 levels over this locus were close to background in wild-type cells and only increased in the absence of the counteracting demethylase Epe1. Our analysis failed to detect H3K9me2 over *ncRNA.394* in untreated wild-type cells (Fig. 2b and Extended Data Fig. 3a).

Deletion of *ncRNA.394* did not result in caffeine resistance (Extended Data Fig. 3b). Prolonged non-selective growth without caffeine of cells exhibiting the *ncRNA.394* H3K9me2 domain resulted in loss of H3K9me2 over this region, whereas growth with caffeine present extended the H3K9me2 domain upstream to include the *SPBC17G9.13c*^+^ and *SPBC17G9.12c*^+^ genes (Extended Data Fig. 3c). Deletion of *SPBC17G9.12c*^+^ or *eno101*^+^ did not result in caffeine resistance (Extended Data Fig. 3b). *SPBC17G9.13c*+ is essential for viability precluding testing a deletion mutant for resistance. Together these analyses suggest that reduced expression of *SPBC17G9.13c*+ may mediate caffeine resistance.

To test directly if heterochromatin formation at these specific loci can result in caffeine resistance, *tetO* DNA binding sites were inserted at the *hba1* and *ncRNA.394* loci and a TetR-Clr4* (catalytically active but lacking the Clr4 chromodomain) fusion protein expressed to force assembly of synthetic heterochromatin upon recruitment to these loci^4,5^. Combining *tetO* with TetR-Clr4* in the absence of anhydrotetracycline (-AHT) resulted in a novel H3K9me2 domain at each locus and growth of cells in the presence of caffeine (Fig. 3 and Extended Data Fig. 4). This indicates that heterochromatin-mediated silencing at either the *hba1* or *ncRNA.394* loci results in caffeine resistance. Because TetR-Clr4* tethering close to *SPBC17G9.13c*^+^ resulted in caffeine resistance we surmise that reduced expression of the *SPBC17G9.13c*^+^ gene upstream of *ncRNA.394* is likely responsible for caffeine resistance at this locus.

**Figure 3.**
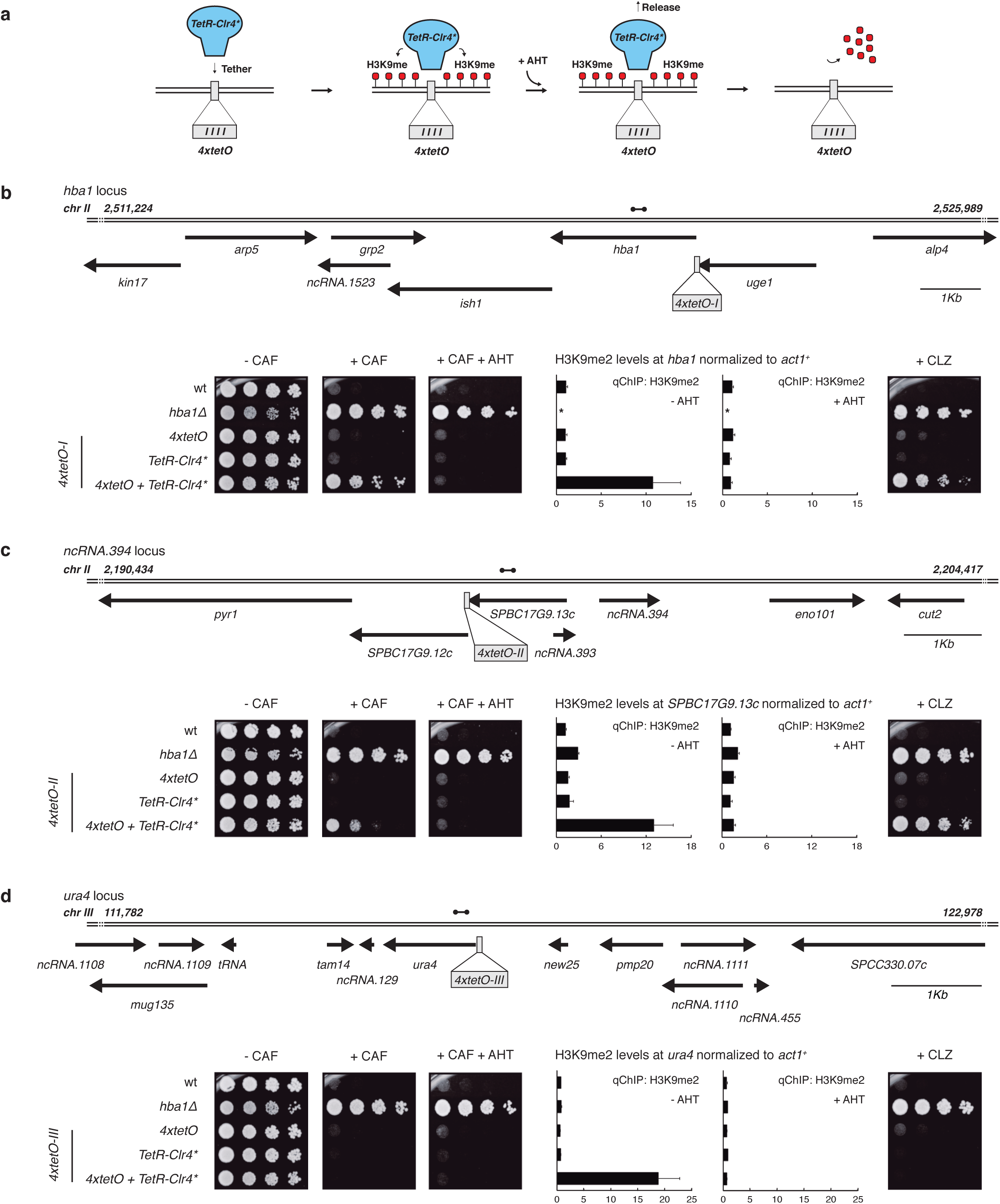
Forced synthetic heterochromatin placement at the identified loci is sufficient to drive caffeine resistance in wild-type cells. **a**, Diagram illustrating *TetR-Clr4\**-mediated H3K9me deposition at *4xtetO* binding sites. Addition of anhydrotetracycline (+AHT) causes release of *TetR-Clr4\** from *4xtetO* sites which results in active removal of H3K9me. **b-d**, Wild-type cells harbouring *4xtetO* binding sites at the *hba1* or *ncRNA.394* loci (or *ura4* as control) and expressing *TetR-Clr4\** were assessed for caffeine resistance in the absence or presence of AHT. Quantitative chromatin immunoprecipitation (qChIP) of H3K9me2 levels on *hba1* (**b**), *SPBC17G9.13c* (**c**) and *ura4* (**d**) loci. Data are mean ± SD (error bars) (n = 3 experimental replicates). Dumbbells indicate oligonucleotides used. *Note *hba1* is not present in *hba1δ*. Strains were also assessed for resistance to the fungicide clotrimazole.

Remarkably, we found that strains with forced synthetic heterochromatin at either *hba1* or *ncRNA.394* loci displayed resistance to the widely-used clinical fungicide clotrimazole (Fig. 3, +CLZ). Further investigation of our unstable caffeine-resistant isolates revealed that those with heterochromatin formation at the *hba1* (UR-1) and the *ncRNA.394* (UR-2) loci are also resistant to clotrimazole and generate small interfering RNAs (siRNAs) homologous to the surrounding genes (Extended Data Fig. 5).

In addition to a heterochromatin domain over *ncRNA.394*, analysis of ChIP-seq input DNA indicated that many independent unstable caffeine-resistant isolates also contained overlapping regions of chromosome III present at increased copy number (Extended Data Fig. 6). In 11 of 12 isolates, the minimal region of overlap contains the *cds1*^+^ gene, overexpression of which is known to confer caffeine resistance^17^. To determine if amplification of the *cds1* locus occurred before or after formation of the *ncRNA.394* H3K9me2 domain we analyzed a sample frozen later in the time series for the same isolate (UR-2). ChIP-seq analysis showed that the *ncRNA.394* H3K9me2 domain was present in the initial caffeine-resistant isolate (4 days +CAF), whereas the *cds1* locus amplification arose later (7 days +CAF) (Extended Data Fig. 7a). These data suggest that development of resistance is a multistep process in which a combination of different events can increase resistance. In agreement with this hypothesis, deletion of *clr4*^+^ in the initial UR-2 isolate (4 days +CAF) resulted in loss of caffeine resistance in all transformants tested (6/6) (Extended Data Fig. 7b and 1c). However, only half of the transformants (3/6) lost resistance to caffeine when *clr4*^+^ was deleted in the isolate displaying *cds1* locus amplification (7 days +CAF), suggesting that once amplification of the *cds1* locus occurs heterochromatin is not required for resistance. In UR-2 a new heterochromatin domain occurred before *cds1*^+^ amplification but it is possible that events are stochastic and occur in no fixed order. Interestingly, both events – the *ncRNA.394* H3K9me2 domain and *cds1* locus amplification – are unstable and lost following growth in the absence of caffeine (Extended Data Fig. 7c).

To investigate the dynamics of heterochromatin domain formation in response to caffeine we exposed wild-type cells to low (7 mM) or medium (14 mM) doses of caffeine for 18 hours. Cells in low caffeine accomplished ∼8 doublings, whereas fewer than 3 population doublings occurred in medium caffeine. ChIP-seq for H3K9me2 identified several new ectopic domains of heterochromatin following exposure to low caffeine. Ectopic domains were detected at loci known to accumulate H3K9me2 in the absence of Epe1^8^, including *ncRNA.394* (Fig. 4a, top). Remarkably, following treatment with medium doses of caffeine, ectopic heterochromatin was restricted to *ncRNA.394*, and H3K9me2 levels at this locus were approximately 2-fold greater than those after exposure to low caffeine (Fig. 4a, bottom). Together these data indicate that, when exposed to near-lethal doses of caffeine (medium, 14 mM), wild-type cells can rapidly develop resistance by forming heterochromatin over a locus (*ncRNA.394*) that confers resistance when silenced.

**Figure 4.**
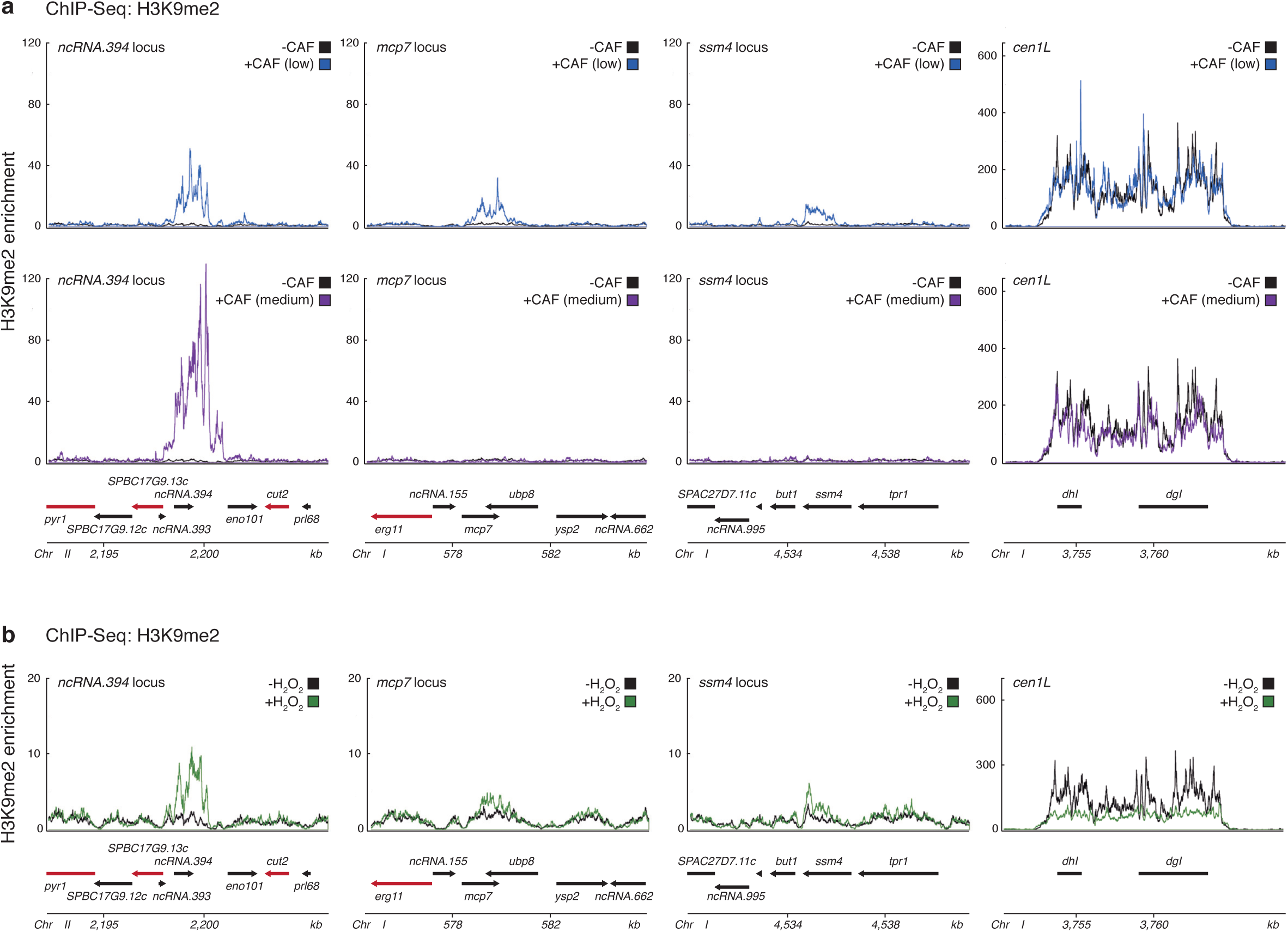
Dynamic heterochromatin redistribution following short exposure to external insults in wild-type cells. **a**, H3K9me2 ChIP-seq enrichment at *ncRNA.394, mcp7* and *ssm4* loci following 18 hr exposure to low (7 mM, top) or medium (14 mM, bottom) concentrations of caffeine. **b**, H3K9me2 ChIP-seq enrichment at *ncRNA.394, mcp7* and *ssm4* loci following 18hr exposure to a low concentration of hydrogen peroxide (1 mM). **a-b**, Data are represented as relative fold enrichment over input and compared to levels in wt cells. Relevant genes within and flanking ectopic heterochromatin domains are indicated. Red arrows indicate essential genes. H3K9me2 enrichment at pericentromeric *dhI* and *dgI* repeats (*cen1L*) of chromosome I shown as control (note different scale).

To determine if other insults also induce novel heterochromatin domains, we exposed wild-type cells to oxidative stress by addition of hydrogen peroxide (1 mM). ChIP-seq for H3K9me2 revealed the presence of ectopic heterochromatin domains at similar locations to those observed in low caffeine treatment, albeit H3K9me2 levels were lower (Fig. 4b). Thus, our results reveal an adaptive epigenetic response following exposure to external insults, and suggest that stress-response pathways may regulate activities that modulate heterochromatin formation thereby ensuring cell survival in fluctuating environmental conditions (Extended Data Fig. 8).

It is well known that DNA methylation-dependent epimutations arise in plants and are propagated by maintenance methyltransferases^18,19^. RNAi-mediated epimutations have been shown to arise in the fungus *Mucor circinelloides*^20^, but it is not known if these are DNA methylation or heterochromatin dependent. As fission yeast lacks DNA methylation^21,22^ this epigenetic mark cannot be responsible for the epimutations described here. Instead our analyses indicate that these adaptive epimutations are transmitted in wild-type cells by the previously-identified Clr4/H3K9me read-write mechanism^4,5^.

Our findings prompt the question as to why epimutants have not been detected previously in mutant screens. Phenotypic screens are usually very stringent, and generally only the strongest mutants are retained for further investigation and eccentric mutants are discarded. Here we essentially select for weak mutants by applying low doses of a drug that is at the threshold of preventing the growth of most cells. Selection was applied for a short period of time in order to maximize the chance of identifying isolates that exhibit unstable phenotypes prior to the development of genetic alterations.

Fungal infections are on the rise, especially in immunocompromised humans. There are few effective anti-fungal agents and resistance is rendering them increasingly ineffective^23,24^. The widespread use of related azole compounds to control fungal deterioration of crops may leave low fungicide levels in the soil, possibly leading to the unwitting selection of resistant epimutants in fungi, similar to those described here, that may ultimately drive the increasing number of cases of azole-resistant Aspergillosis and Cryptococcosis in the clinic. Use of the existing battery of so called ‘epigenetic drugs’ - compounds that inhibit histone modifying enzymes - may identify molecules that block heterochromatin formation and hence reduce the emergence of anti-fungal resistance in plant and animal pathogenic fungi.

## Methods

### Yeast strains and manipulations

Standard methods were used for fission yeast growth, genetics and manipulation^25^. *S. pombe* strains used in this study are described in Supplementary Information Table S2. Oligonucleotide sequences are listed in Supplementary Information Table S3. For pDUAL-adh21-TetR-2xFLAG-Clr4-CDΔ (abbreviated as TetR-Clr4*), the *nmt81* promoter of pDUAL-nmt81-TetR-2xFLAG-Clr4-CDΔ^4^, was replaced by the *adh21* promoter (pRAD21, gift from Y. Watanabe). *Not*I-digested plasmid was integrated at *leu1*^+^. Pap1-N424STOP strain and strains carrying *4xtetO* insertions were constructed by CRISPR/Cas9-mediated genome editing using the *SpEDIT* system (Allshire Lab; available on request) with oligonucleotides listed in Supplementary Information Table S3. Yeast extract plus supplements (YES) was used to grow all cultures. 16 mM caffeine (Sigma, C0750) was added to media for caffeine resistance screens and serial dilution assays. Caffeine-resistant colonies that formed after seven days were picked and patched to +CAF plates. After four days of growth, isolates were frozen (4 days +CAF). 4 days +CAF isolates were repatched and grown for three days on +CAF plates and then frozen (7 days +CAF). Subsequently, 7 days +CAF isolates were repatched every three days on +CAF plates up to twenty days of total growth on +CAF plates (20 days +CAF). 0.29 μM clotrimazole (Sigma, C6019) was added to media for clotrimazole resistance serial dilution assays. 7 or 14 mM caffeine (Sigma, C0750), or 1 mM hydrogen peroxide (Sigma, H1009) were added to media for 18 hours for drug treatment experiments. To release TetR-Clr4*, 10 μM anhydrotetracycline (AHT) was added to the media.

### Serial dilution assays

Equal amounts of starting cells were serially diluted four-fold and then spotted onto appropriate media. Cells were grown at 30-32°C for 3-5 days and then photographed.

### Chromatin immunoprecipitation (ChIP)

ChIP experiments were performed as previously described^26^ using anti-H3K9me2 (5.1.1, a kind gift by Takeshi Urano). Immunoprecipitated DNA was recovered with Chelex-100 resin (BioRad) for ChIP-qPCR (qChIP) experiments or with QIAquick PCR Purification Kit (Qiagen) for ChIP-seq experiments.

### Quantitative ChIP (qChIP)

qChIPs were analysed by real-time PCR using Lightcycler 480 SYBR Green (Roche) with oligonucleotides listed in Supplementary Information Table S3. All ChIP enrichments were calculated as % DNA immunoprecipitated at the locus of interest relative to the corresponding input samples and normalized to % DNA immunoprecipitated at the *act1*^+^ locus. Histograms represent data averaged over three biological replicates. Error bars represent standard deviations.

### ChIP-seq library preparation and analysis

Illumina-compatible libraries were prepared as previously described^26^ using NEXTflex-96 barcode adapters (Bioo Scientific) and Ampure XP beads (Beckman Coulter). Libraries were then pooled to allow multiplexing and sequenced on an Illumina HiSeq2000, NextSeq or MiniSeq system (150-cycle high output kit) by 75 bp paired-end sequencing.

Approximately 6-10 million 75 bp paired-end reads were produced for each sample. Raw reads were then de-multiplexed and trimmed using Trimmomatic (v0.35)^27^ to remove adapter contamination and regions of poor sequencing quality. Trimmed reads were aligned to the *S. pombe* reference genome (*972h*^*-*^, ASM294v2.20) using Bowtie2 (v2.3.3)^28^. Resulting bam files were processed using Samtools (v1.3.1)^29^ and picard-tools (v2.1.0) (http://broadinstitute.github.io/picard) for sorting, removing duplicates and indexing. Coverage bigwig files were generated by BamCoverage (deepTools v2.0) and ratios IP/input were calculated using BamCompare (deepTools v2.0)^30^ in SES mode for normalisation^31^. Peaks were called using MACS2^32^ in PE mode and broad peak calling (broad-cutoff = 0.05). Region-specific H3K9me2 enrichment plots were generated using the Sushi R package (v1.22)^33^.

### SNP and indel calling

SNPs and indels were called as described^34^. Trimmed reads were mapped to the *S. pombe* reference genome (*972h*^*-*^, ASM294v2.20) using Bowtie2 (v2.3.3)^28^. GATK^35,36^ was used for base quality score recalibration. SNPs and indels were called with GATK HaplotypeCaller^35,36^ and filtered using custom parameters. Functional effect of variants was determined using Variant Effect Predictor^37^.

### Copy number variation analysis

Copy number variation was determined using CNVkit^38^ in Whole-Genome Sequencing (-wgs) mode. Wild-type ChIP-seq input bam files were used as reference.

### qRT–PCR analysis

For qRT-PCR, total RNA was extracted using the Monarch Total RNA Miniprep Kit (New England Biolabs) according to the manufacturer’s instructions. Contaminating DNA was removed by treating with Turbo DNase (Invitrogen) and reverse transcription was performed using LunaScript RT Supermix Kit (New England Biolabs). Oligonucleotides used for qRT-PCR are listed in Supplementary Information Table S3. qRT-PCR histograms represent three biological replicates; error bars correspond to the standard deviation. * *P* < 0.05 (*t* test).

### Small RNA-seq

50 mL of log-phase cells were collected and processed using the mirVana miRNA Isolation kit (Invitrogen). Resulting sRNA was treated with TURBO DNase (Invitrogen) and used for library construction using NEBNext Multiplex Small RNA Library Prep Set for Illumina (New England Biolabs) according to manufacturer’s instructions. Libraries were pooled and sequenced on an Illumina NextSeq platform by 50 bp single-end sequencing. Raw reads were then de-multiplexed and processed using Cutadapt (v1.17) to remove adapter contamination and discard reads shorter than 19 nucleotides or longer than 25 nucleotides. Coverage plots were generated using SCRAM^39^.

## Acknowledgments

We thank Lorenza Di Pompeo and Andreas Fellas for laboratory support, Pin Tong and Ryan Ard for sharing technical expertise and members of the Allshire lab for valuable discussions. We are grateful to Adrian Bird, Wendy Bickmore and Lucia Massari for comments on the manuscript. We thank Takeshi Urano for kindly providing the 5.1.1 (H3K9me) antibody and Yoshinori Watanabe for the pRAD21 plasmid. S.T-G. was supported by the Darwin Trust of Edinburgh. R.C.A. is a Wellcome Principal Research Fellow (095021, 200885); the Wellcome Centre for Cell Biology is supported by core funding from Wellcome (203149).

## Author contributions

S.T-G., P.N.C.B.A. and R.C.A. conceived the project. S.T-G. and P.N.C.B.A. performed preliminary studies. S.T-G. performed experiments and bioinformatics. M.S. and A.L.P. contributed to ChIP-seq and qChIP experiments. S.A.W. contributed to sRNA-seq experiments. S.T-G., A.L.P. and R.C.A. wrote the manuscript.

---

**Supplementary Information Table 1.** Summary of epigenetic (H3K9me2 domains) and genetic (SNPs, indels and copy number variation) changes found in unstable (UR) caffeine-resistant isolates.

| Isolate | Ectopic heterochromatin location |  | SNPs or indels in coding sequences? | Partial duplication of *Chr III*? |
| --- | --- | --- | --- | --- |
|  | *ncRNA.394* | other loci |  |  |
| UR-1 |  | ✓ (*hba1*) | Clr5-Q264STOP / Meu27-S100Y |  |
| UR-2 | ✓ |  | Sdo1-R11C |  |
| UR-3 |  | ✓ (*ppr4*) | Clr5-Q264STOP / Meu27-S100Y |  |
| UR-4 |  | ✓ (*grt1*) | - | ✓ |
| UR-5 |  | ✓ (*fio1*) | Clr5-Q264STOP / Meu27-S100Y | ✓ |
| UR-6 |  | ✓ (*mbx2*) | - | ✓ |
| UR-7 |  | ✓ (*ppr4*) | Clr5-Q264STOP / Meu27-S100Y |  |
| UR-8 | ✓ |  | - |  |
| UR-9 | ✓ |  | - |  |
| UR-10 | ✓ |  | Cob1-F318L |  |
| UR-11 | ✓ |  | - |  |
| UR-12 | ✓ |  | - |  |
| UR-13 | ✓ |  | - | ✓ |
| UR-14 | ✓ |  | Npp-W300STOP / SPBC16H5.13-S1011L | ✓ |
| UR-15 | ✓ |  | - | ✓ |
| UR-16 | ✓ |  | - |  |
| UR-17 | ✓ |  | SPCC777.02-R120R | ✓ |
| UR-18 | ✓ |  | SPCC777.02-R120R | ✓ |
| UR-19 | ✓ |  | Sdo1-R11C | ✓ |
| UR-20 | ✓ |  | - |  |
| UR-21 | ✓ |  | - | ✓ |
| UR-22 | ✓ |  | - | ✓ |
| UR-23 | ✓ |  | Pch1-Q234STOP |  |
| UR-24 | ✓ |  | - |  |
| UR-25 | ✓ |  | - | ✓ |
| UR-26 | ✓ |  | SPBC1271.08c-A133A |  |
| UR-27 | ✓ |  | SPCC4B3.13-A229V |  |
| UR-28 | ✓ |  | Mug72-N116S |  |
| UR-29 | ✓ |  | Mug72-N116S |  |
| UR-30 | ✓ |  | - |  |

**Supplementary Information Table 2.** *Schizosaccharomyces pombe* strains used in this study.

| Strain number | Name | Description |
| --- | --- | --- |
| 143 | wt | *h*- ED972 wild-type |
| B4411 | SR-1 | Stable 16 mM Caffeine Resistant Isolate – From wt – 1 |
| B4412 | SR-2 | Stable 16 mM Caffeine Resistant Isolate – From wt – 2 |
| B4413 | UR-1 | Unstable 16 mM Caffeine Resistant Isolate – From wt – 1 |
| B4414 | UR-2 | Unstable 16 mM Caffeine Resistant Isolate – From wt – 2 |
| B4415 | UR-3 | Unstable 16 mM Caffeine Resistant Isolate – From wt – 3 |
| B4416 | UR-4 | Unstable 16 mM Caffeine Resistant Isolate – From wt – 4 |
| B4417 | UR-5 | Unstable 16 mM Caffeine Resistant Isolate – From wt – 5 |
| B4418 | UR-6 | Unstable 16 mM Caffeine Resistant Isolate – From wt – 6 |
| B4419 | UR-7 | Unstable 16 mM Caffeine Resistant Isolate – From wt – 7 |
| B4420 | UR-8 | Unstable 16 mM Caffeine Resistant Isolate – From wt – 8 |
| B4421 | UR-9 | Unstable 16 mM Caffeine Resistant Isolate – From wt – 9 |
| B4422 | UR-10 | Unstable 16 mM Caffeine Resistant Isolate – From wt – 10 |
| B4423 | UR-11 | Unstable 16 mM Caffeine Resistant Isolate – From wt – 11 |
| B4424 | UR-12 | Unstable 16 mM Caffeine Resistant Isolate – From wt – 12 |
| B4425 | UR-13 | Unstable 16 mM Caffeine Resistant Isolate – From wt – 13 |
| B4426 | UR-14 | Unstable 16 mM Caffeine Resistant Isolate – From wt – 14 |
| B4427 | UR-15 | Unstable 16 mM Caffeine Resistant Isolate – From wt – 15 |
| B4428 | UR-16 | Unstable 16 mM Caffeine Resistant Isolate – From wt – 16 |
| B4429 | UR-17 | Unstable 16 mM Caffeine Resistant Isolate – From wt – 17 |
| B4430 | UR-18 | Unstable 16 mM Caffeine Resistant Isolate – From wt – 18 |
| B4431 | UR-19 | Unstable 16 mM Caffeine Resistant Isolate – From wt – 19 |
| B4432 | UR-20 | Unstable 16 mM Caffeine Resistant Isolate – From wt – 20 |
| B4433 | UR-21 | Unstable 16 mM Caffeine Resistant Isolate – From wt – 21 |
| B4434 | UR-22 | Unstable 16 mM Caffeine Resistant Isolate – From wt – 22 |
| B4435 | UR-23 | Unstable 16 mM Caffeine Resistant Isolate – From wt – 23 |
| B4436 | UR-24 | Unstable 16 mM Caffeine Resistant Isolate – From wt – 24 |
| B4437 | UR-25 | Unstable 16 mM Caffeine Resistant Isolate – From wt – 25 |
| B4438 | UR-26 | Unstable 16 mM Caffeine Resistant Isolate – From wt – 26 |
| B4439 | UR-27 | Unstable 16 mM Caffeine Resistant Isolate – From wt – 27 |
| B4440 | UR-28 | Unstable 16 mM Caffeine Resistant Isolate – From wt – 28 |
| B4441 | UR-29 | Unstable 16 mM Caffeine Resistant Isolate – From wt – 29 |
| B4442 | UR-30 | Unstable 16 mM Caffeine Resistant Isolate – From wt – 30 |
| B4443 | SR-1 *clr4Δ* - 1 | SR-1 *clr4Δ::NAT* - transformant 1 |
| B4444 | SR-1 *clr4Δ* - 2 | SR-1 *clr4Δ::NAT* - transformant 2 |
| B4445 | SR-1 NAT control - 1 | SR-1 *NAT:3' of ura4* - transformant 1 |
| B4446 | SR-1 NAT control - 2 | SR-1 *NAT:3' of ura4* - transformant 2 |
| B4447 | SR-2 *clr4Δ* - 1 | SR-2 *clr4Δ::NAT* - transformant 1 |
| B4448 | SR-2 *clr4Δ* - 2 | SR-2 *clr4Δ::NAT* - transformant 2 |
| B4449 | SR-2 NAT control - 1 | SR-2 *NAT:3' of ura4* - transformant 1 |
| B4450 | SR-2 NAT control - 2 | SR-2 *NAT:3' of ura4* - transformant 2 |
| B4451 | UR-1 *clr4Δ* - 1 | UR-1 *clr4Δ::NAT* - transformant 1 |
| B4452 | UR-1 *clr4Δ* - 2 | UR-1 *clr4Δ::NAT* - transformant 2 |
| B4453 | UR-1 NAT control-1 | UR-1 *NAT:3' of ura4* - transformant 1 |
| B4454 | UR-1 NAT control-2 | UR-1 *NAT:3' of ura4* - transformant 2 |
| B4455 | UR-2 *clr4Δ* - 1 | UR-2 *clr4Δ::NAT* - transformant 1 |
| B4456 | UR-2 *clr4Δ* - 2 | UR-2 *clr4Δ::NAT* - transformant 2 |
| B4457 | UR-2 NAT control - 1 | UR-2 *NAT:3' of ura4* - transformant 1 |
| B4458 | UR-2 NAT control - 2 | UR-2 *NAT:3' of ura4* - transformant 2 |
| B4352 | Pap1-N424STOP | *h*- *pap1-N424STOP* |
| B4459 | UR-2 +14 days -CAF | UR-2 after growth on -CAF media for 14 days |
| B4460 | *hba1Δ* | *h*- *hba1Δ::NAT* |
| B4461 | *SPBC17G9.12cΔ* | *h*- *SPBC17G9.12cΔ::NAT* |
| B4462 | *ncRNA.393Δ* | *h*- *ncRNA.393Δ::NAT* |
| B4463 | *ncRNA.394Δ* | *h- ncRNA.394Δ::NAT* |
| B4464 | *eno101Δ* | *h- eno101Δ::NAT* |
| B3797 | *TetR-Clr4\** | *h+ leu1+adh21-TetROFF-2xFLAG-Clr4-cdd* |
| B3808 | *4xtetO-II* | *h- 4xtetO 3' of SPBC17G9.13c leu1-32* |
| B3813 | *4xtetO-I* | *h- 4xtetO 5' of hba1 leu1-32* |
| B3820 | *4xtetO-III* | *h- 4xtetO 5' of ura4 leu1-32* |
| B4465 | *TetR-Clr4\* + 4xtetO-II* | *h+ leu1+adh21-TetROFF-2xFLAG-Clr4-cdd 4xtetO 3' of SPBC17G9.13c* |
| B4466 | *TetR-Clr4\* + 4xtetO-I* | *h+ leu1+adh21-TetROFF-2xFLAG-Clr4-cdd 4xtetO 5' of hba1* |
| B4467 | *TetR-Clr4\* + 4xtetO-III* | *h+ leu1+adh21-TetROFF-2xFLAG-Clr4-cdd 4xtetO 5' of ura4* |
| B4468 | UR-2 (7 days +CAF) | UR-2 after growth on +CAF media for 3 days |
| B4469 | UR-2 (7 days +CAF<br>→14 days -CAF) | UR-2 after growth on +CAF media for 3 days and then on -CAF media for 14 days |

**Supplementary Information Table 3.**
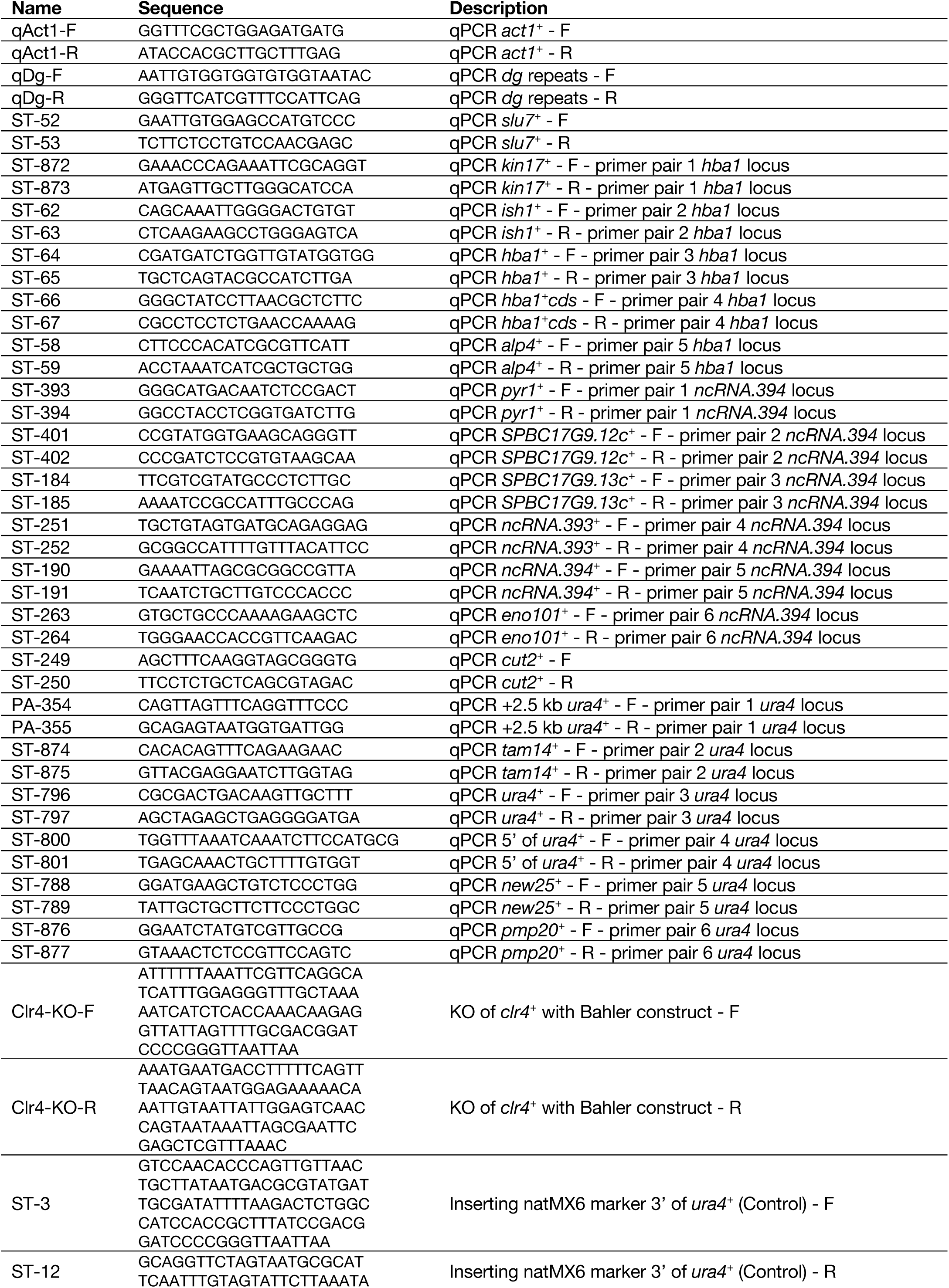
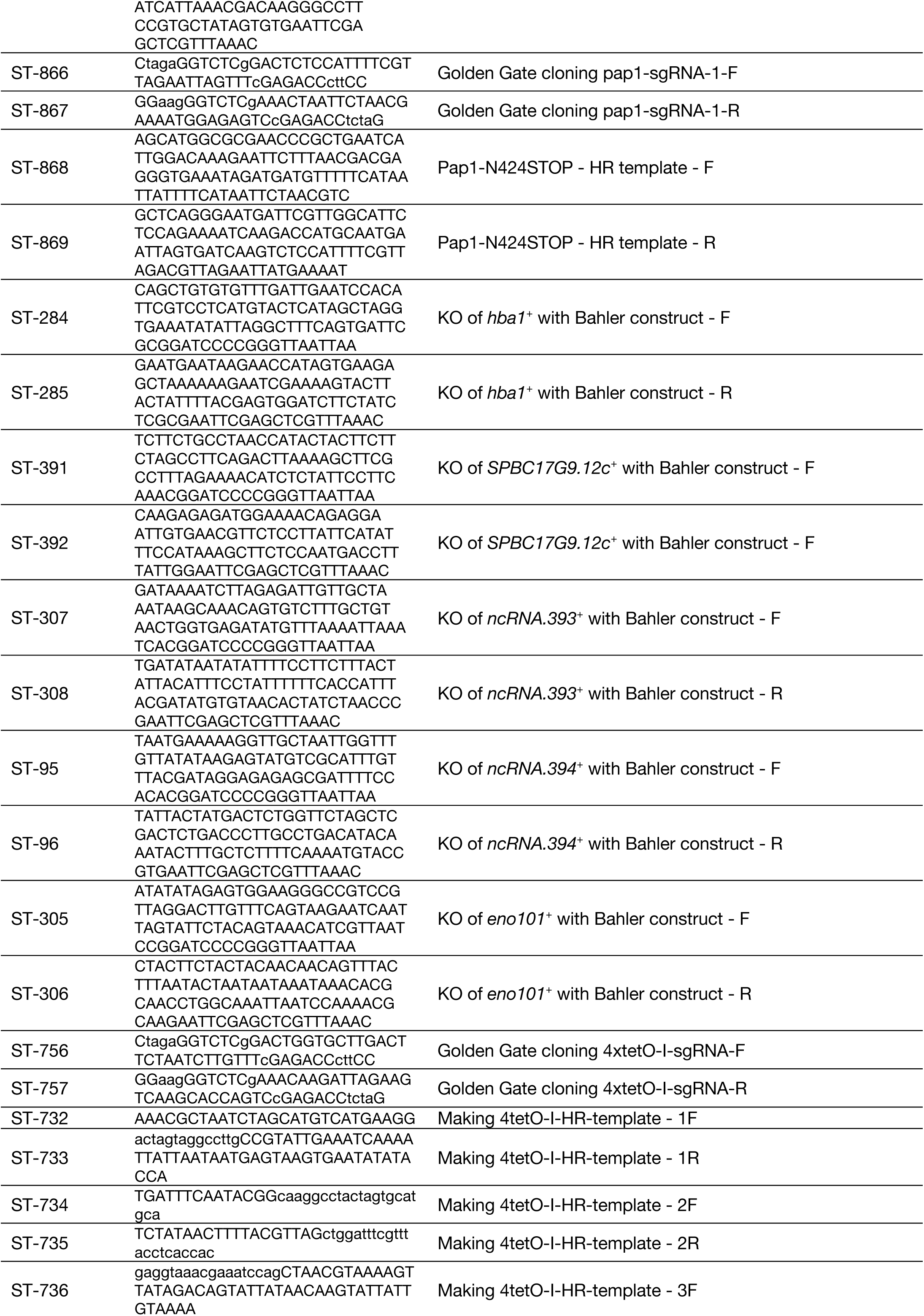
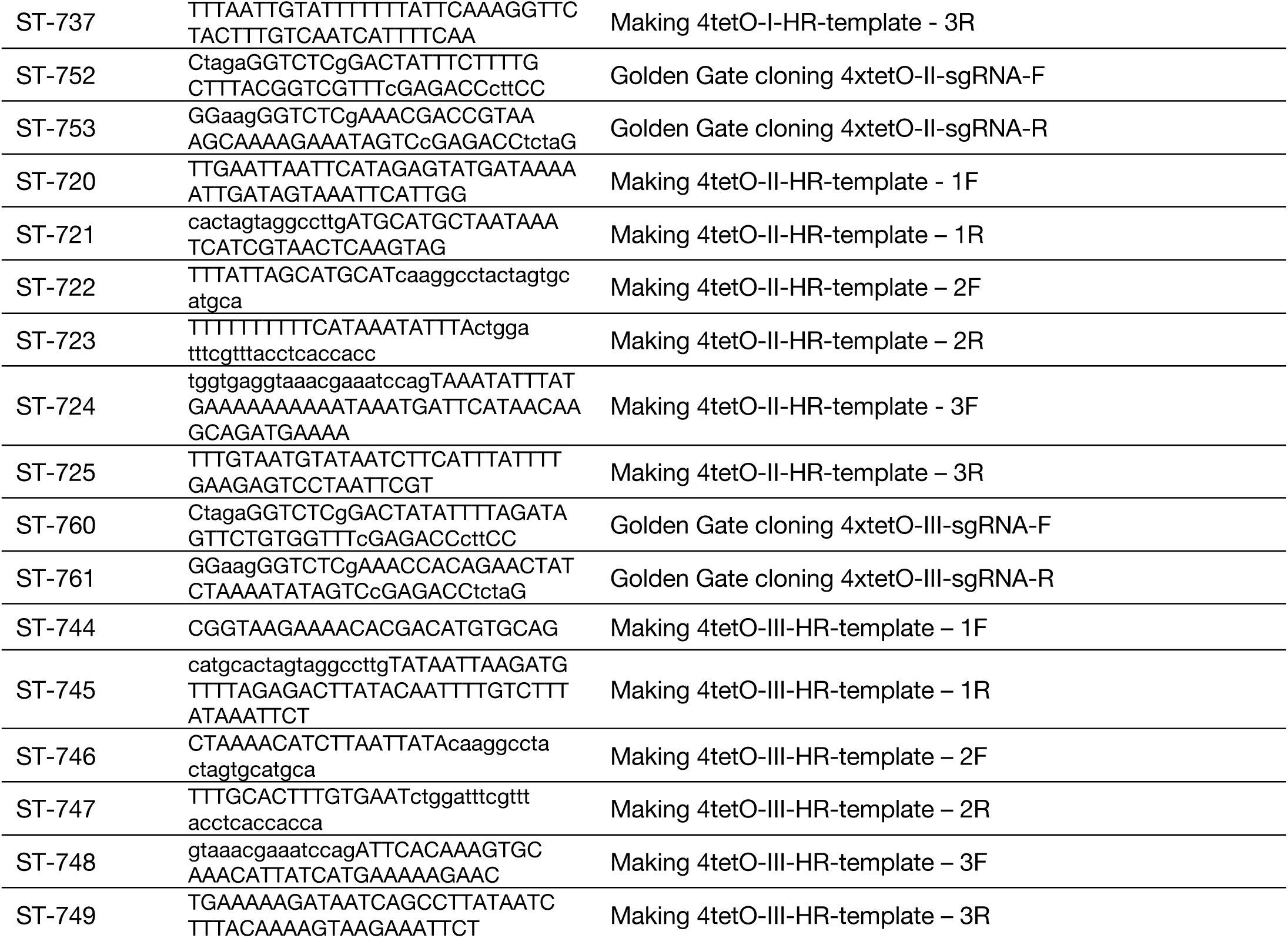
Oligonucleotides used in this study.

**Extended Data Figure 1.**
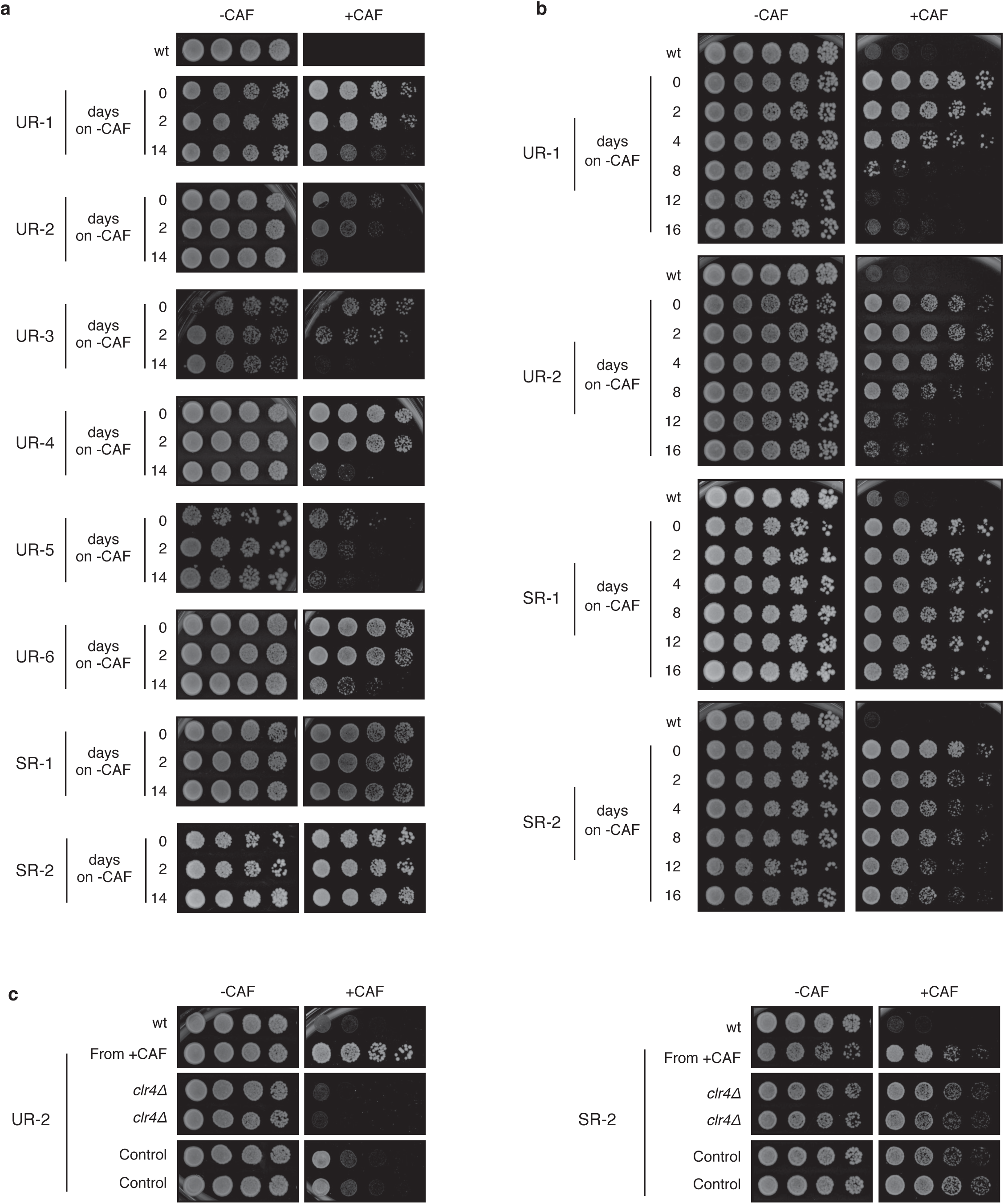
Identification of heterochromatin-dependent epimutants resistant to caffeine. **a**, Unstable (UR) and stable (SR) caffeine-resistant isolates were identified using our screening strategy. After growth on non-selective media for 14 days caffeine resistance is lost in UR isolates but not in SR isolates. **b**, Caffeine resistance is lost progressively in unstable (UR) isolates but maintained in stable (SR) isolates. **c**, Caffeine resistance in UR isolates depends on the Clr4 H3K9 methyltransferase. *clr4*+ (*clr4δ*) or an unlinked intergenic region (Control) were deleted in unstable (UR-2) and stable (SR-2) caffeine-resistant isolates.

**Extended Data Figure 2.**
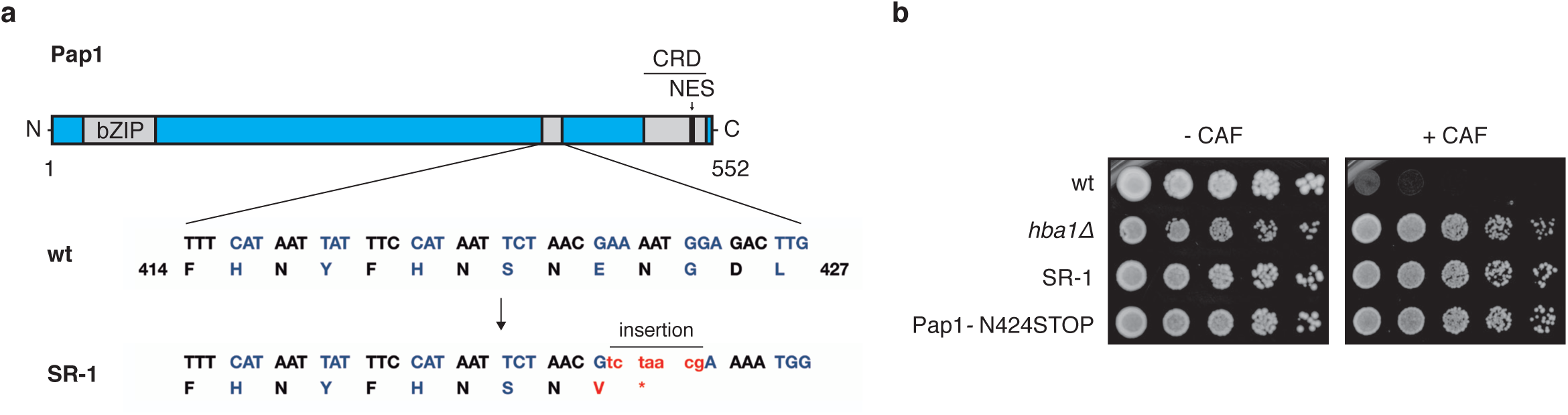
A mutation in *pap1*+ confers caffeine resistance in the stable isolate SR-1. **a**, High-throughput sequencing of the stable isolate SR-1 revealed a 7-nucleotide insertion in *pap1*+. The insertion results in a truncated version of Pap1 (Pap1-N424STOP) lacking the Nuclear Export Signal (NES). **b**, Pap1-N424STOP is resistant to caffeine. The 7-nucleotide insertion identified in SR-1 was introduced in wt cells (Pap1-N424STOP) and caffeine resistance was assessed. *hba1δ* and SR-1 cells were used as positive controls.

**Extended Data Figure 3.**
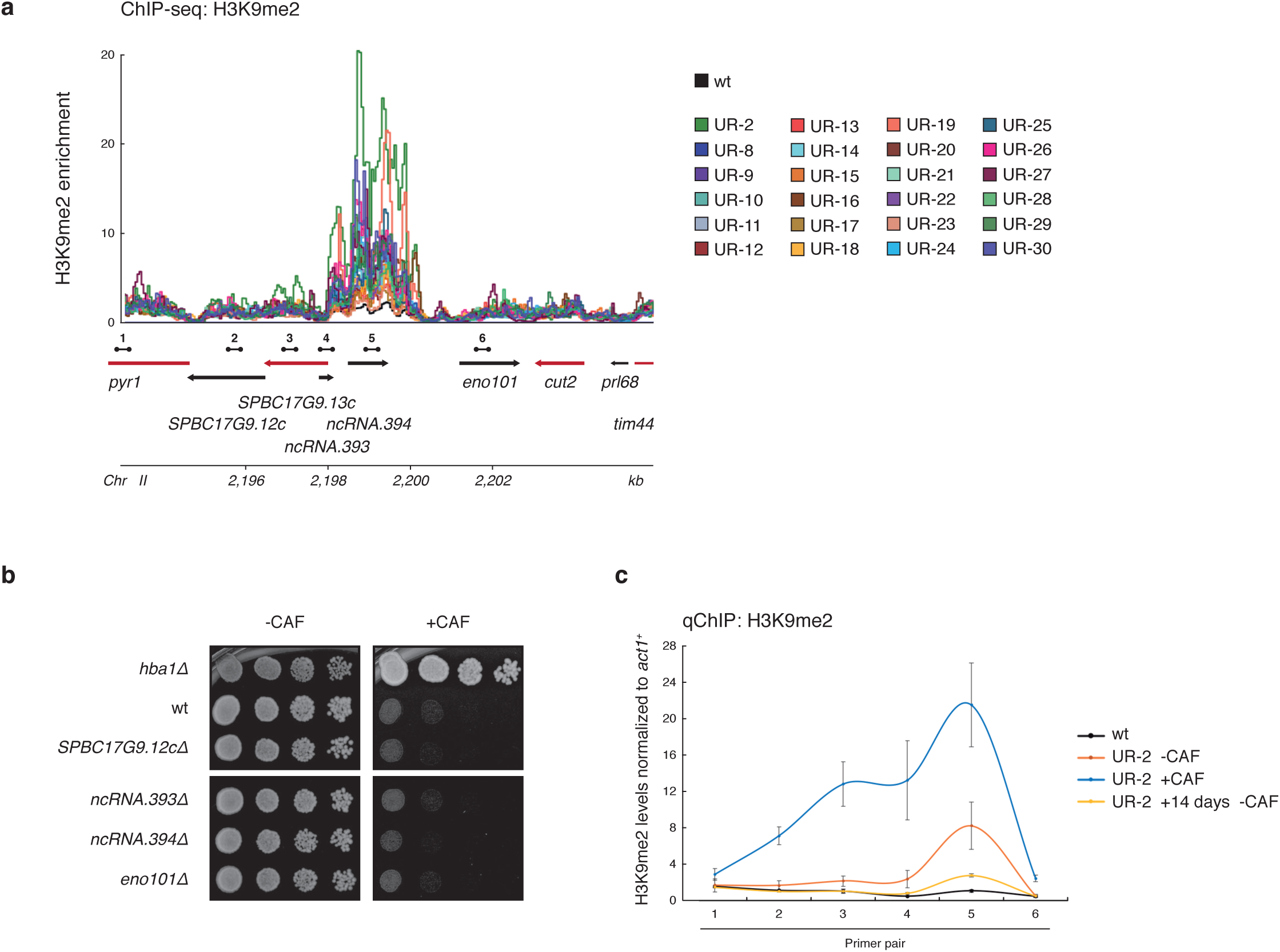
24 of 30 unstable (UR) caffeine-resistant isolates present an ectopic heterochromatin domain over the *ncRNA.394* locus. **a**, H3K9me2 ChIP-seq enrichment at the *ncRNA.394* locus in individual isolates. Data are represented as relative fold enrichment over input and compared to levels in wt cells. Relevant genes within and flanking ectopic heterochromatin domains are indicated. Red arrows indicate essential genes. Dumbbells indicate oligonucleotides used in **c**. **b**, Deletion of *ncRNA.394* or non-essential adjacent genes does not result in caffeine resistance. **c**, Quantitative chromatin immunoprecipitation (qChIP) of H3K9me2 levels at the *ncRNA.394* locus in UR-2 cells. UR-2 cells were grown in the absence (-CAF) or presence (+CAF) of caffeine overnight or in the absence of caffeine for 14 days (+14 days -CAF). Data are mean ± SD (error bars) (n = 3 experimental replicates). Oligonucleotides used are indicated in **a**.

**Extended Data Figure 4.**
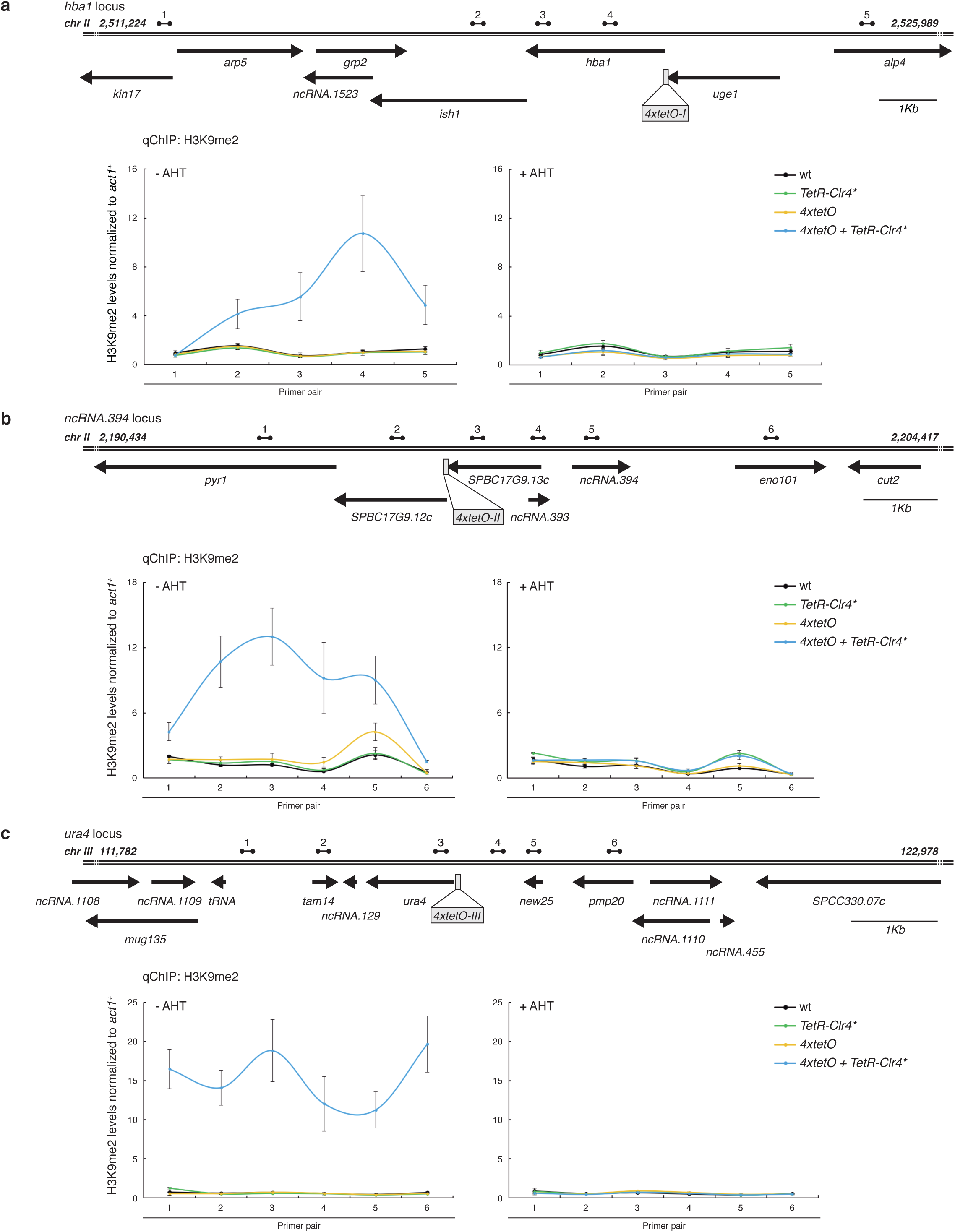
Forced synthetic heterochromatin placement at the identified loci is sufficient to drive caffeine resistance in wild-type cells. **a-c**, Quantitative chromatin immunoprecipitation (qChIP) of H3K9me2 levels in wild-type cells harbouring *4xtetO* binding sites at the identified ectopic heterochromatin loci (or *ura4* as control) and expressing *TetR-Clr4\** in the absence or presence of AHT. **a**, *hba1* locus. **b**, *ncRNA.394* locus. **c**, *ura4* locus. Data are mean ± SD (error bars) (n = 3 experimental replicates). Dumbbells indicate oligonucleotides used.

**Extended Data Figure 5.**
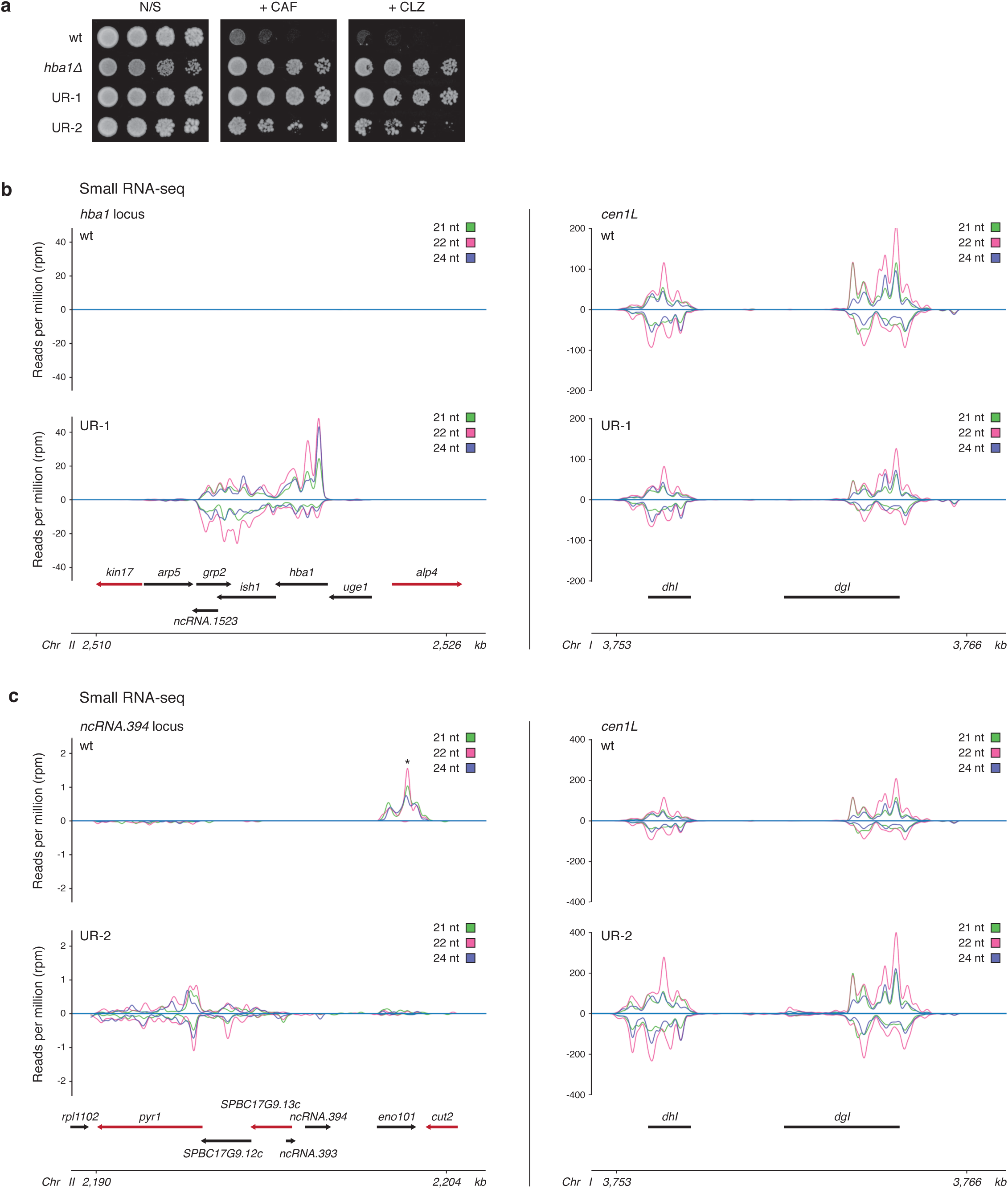
Unstable (UR) caffeine-resistant isolates show cross-resistance to the fungicide clotrimazole and siRNA generation at ectopic heterochromatin domains. **a**, Unstable caffeine-resistant isolates UR-1 and UR-2 were serially diluted and spotted on non-selective (N/S), +CAF and +CLZ plates to assess resistance to caffeine and clotrimazole. **b-c**, Left, small RNA sequencing showing presence of siRNAs (21-24 nucleotides) at ectopic heterochromatin domains in UR-1 (**b**, *hba1* locus) and UR-2 (**c**, *ncRNA.394* locus) cells compared to wt cells. Right, pericentromeric siRNAs mapping to *dhI* and *dgI* repeats (*cen1L*) of chromosome I shown as control. Experiments were performed twice with similar results. *Transcripts mapping to the highly expressed gene *eno101*+ in euchromatic wt conditions (note these are unidirectional RNAs and not siRNAs).

**Extended Data Figure 6.**
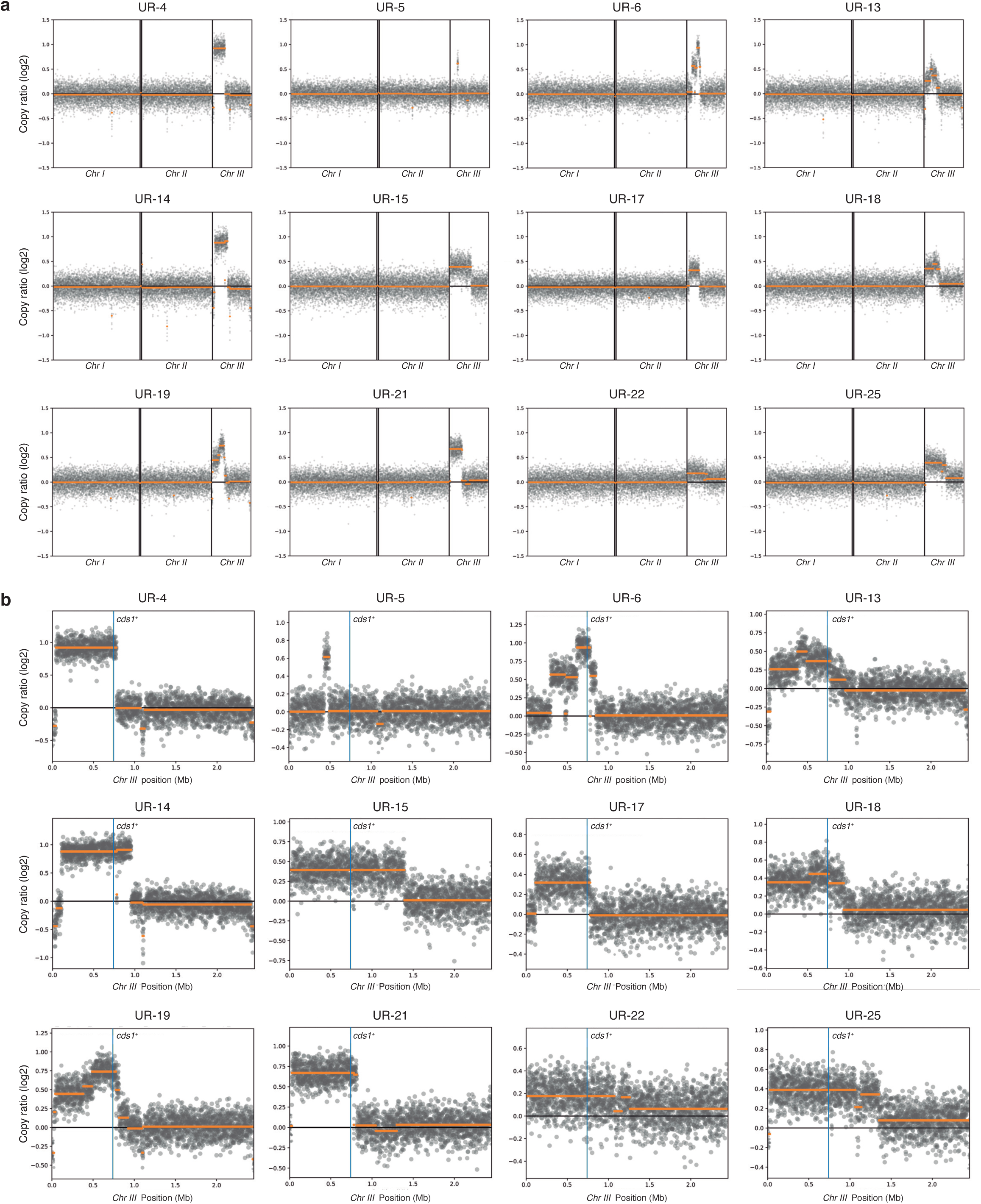
Copy Number Variation (CNV) analysis reveals a partial duplication of chromosome III in 12 of 30 unstable (UR) caffeine-resistant isolates. **a**, Genome-wide coverage plots with overlaid segments in UR isolates showing partial duplication of chromosome III. Wild-type ChIP-seq input data were used as the reference. **b**, Chromosome III coverage plots with overlaid segments in UR isolates showing partial duplication of chromosome III. Location of *cds1*+ is highlighted. Wild-type ChIP-seq input data were used as the reference.

**Extended Data Figure 7.**
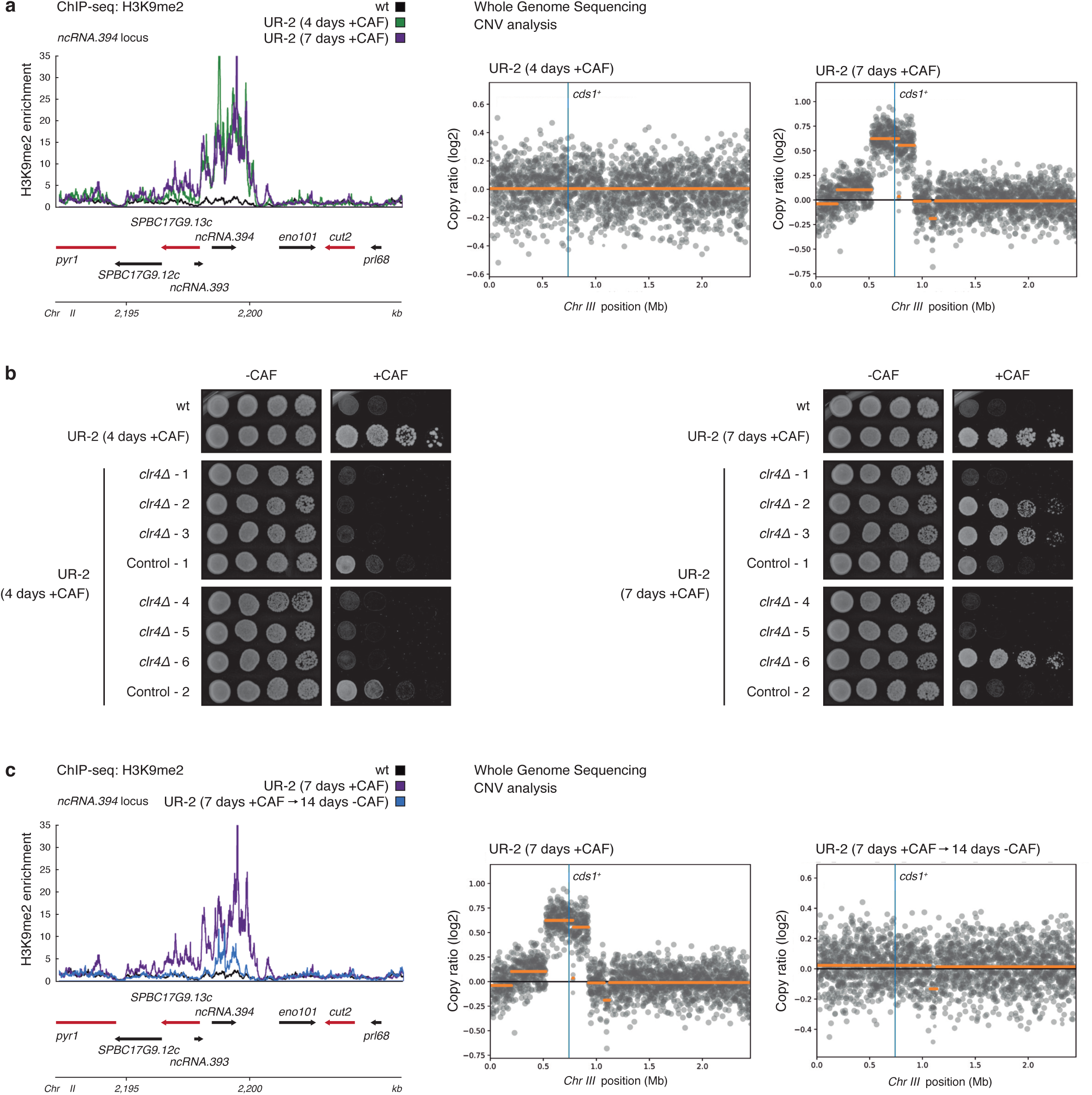
Epigenetic changes preceded genetic changes (CNV) in unstable caffeine-resistant isolate UR-2. **a**, H3K9me2 ChIP-seq enrichment at the *ncRNA.394* locus (left) and chromosome III coverage plots with overlaid segments (right) in UR-2 cells following prolonged growth on +CAF media for 3 days (7 days +CAF). Wild-type ChIP-seq input data were used as the reference for CNV analysis. **b**, *clr4*+ (*clr4δ*) or an unlinked intergenic region (Control) were deleted in UR-2 cells (4 days +CAF) and UR-2 cells after prolonged growth on +CAF media for 3 days (7 days +CAF). All (6/6) UR-2 (4 days +CAF) *clr4δ* transformants lost resistance to caffeine whereas only 50% (3/6) UR-2 (7 days +CAF) lost resistance to caffeine. **c**, H3K9me2 ChIP-seq enrichment at the *ncRNA.394* locus (left) and chromosome III coverage plots with overlaid segments (right) in UR-2 cells following prolonged growth on non-selective media for 14 days after prolonged growth on +CAF media for 3 days (7 days +CAF --> 14 days -CAF). Wild-type ChIP-seq input data were used as the reference for CNV analysis.

**Extended Data Figure 8.**
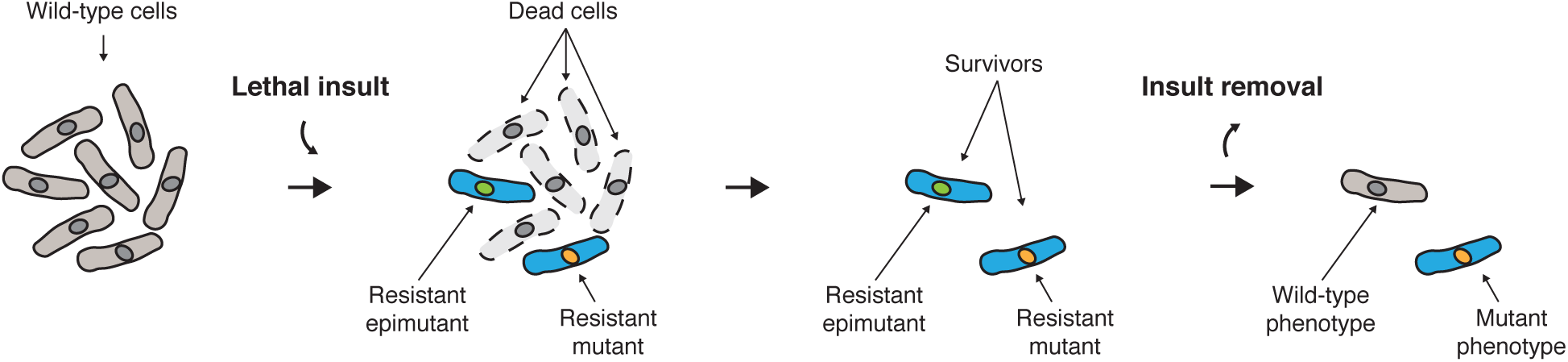
Model. Resistant isolates arise following exposure to a lethal insult. Resistance might be mediated by permanent, DNA-based mutations (resistant mutants) or reversible, heterochromatin-based epimutations (resistant epimutants). Upon insult removal, resistant epimutants can revert to the wild-type phenotype by disassembling ectopic domains of heterochromatin, whereas resistant mutants continue displaying the mutant phenotype due to the genetic nature of DNA mutations.

